# Characterization of α3 Glycine Receptors with Ginkgolide B and Picrotoxin

**DOI:** 10.1101/454710

**Authors:** Sampurna Chakrabarti, Anil Neelakantan, Malcolm M. Slaughter

## Abstract

Ginkgolide B (GB) and picrotoxin (PTX) are antagonists of the major inhibitory receptors of the central nervous system: GABA and glycine receptors (GlyRs). GlyRs contain one or more of the four alpha subunit isoforms of which α1 and α2 have been extensively studied. This report compares GB and PTX block of α3 GlyRs expressed in HEK 293 cells, using whole-cell patch clamp techniques. In CNS, α3 exists as a heteropentamer in conjunction with beta subunits in a 2α:3β ratio. Thus, the nature of block was also tested in α3β heteromeric glycine receptors. GB and PTX blocked α3 GlyRs both in the presence (liganded state) and absence of glycine (unliganded state). This property is unique to α3 subunits; α1 and α2 subunits are only blocked in the liganded state. The GB block of α3 GlyRs is voltage-dependent (more effective when the cell is depolarized) and non-competitive, while the PTX block is competitive and not voltage-dependent. The heteromeric and homomeric α3 GlyRs recovered significantly faster from unliganded GB block compared to liganded GB block, but no such distinction was found for PTX block suggesting more than one binding site for GB. This study sheds light on features of the α3 GlyR that distinguish it from the more widely studied α1 and α2 subunits. Understanding these properties can help decipher the physiological functioning of GlyRs in the CNS and may permit development of subunit specific drugs.

## Introduction

Glycine receptors (GlyRs) and GABA receptors are the major fast inhibitory receptors of the central nervous system. The GlyRs are ionotropic and conduct chloride ions. These cys-loop receptors are pentameric, stoichiometrically made of two α_1-4_ and three β subunits (Durisic et al., 2012). GlyR expression has been found extensively through autoradiography and immunohistochemistry techniques in the central nervous system including in the cerebellar cortex, brainstem, forebrain, retina, and hippocampus (Haverkamp et al., 2003; Crook et al., 2006; Baer et al., 2009; Xu and Gong, 2010). Despite studies on GlyR localization, deciphering the subunit composition of these receptors remains elusive. It is suggested that most GlyRs in native cells are alpha-beta heteromers; α2 homomeric GlyRs expression in fetal rat being the only known exception although there has been speculation that α3 or α1 GlyRs exist in homomeric form in non-somatic locations (Lynch, 2009). The α3β heteromers are often dominant, found in about half of the glycinergic synapses in rat retina and rat dorsal horn of the spinal cord (Haverkamp et al., 2003; Harvey et al., 2004). Relatively few studies have been conducted on α4 GlyRs since it is a pseudo gene in humans (Lynch, 2009) although it’s expression has been shown in rat retina (Heinze et al., 2007). The unique expression of these GlyR subunits in different parts of the CNS suggests that each might have distinct functions. Thus, an important field of research is to develop tools to differentiate between the GlyRs subunits.

Our primary goal in this study was to characterize and compare the blocking action of picrotoxin (PTX) and gingkolide B (GB) on recombinant homomeric and heteromeric α3 GlyRs expressed in human embryonic kidney (HEK 293) cells. GB is a diterpene trilactone extracted from Gingko Biloba leaf (EGb 761). EGb 761 is widely used clinically as a treatment for neurodegeneration. GB, the most potent component of EGb 761, acts as an anti-convulsant, a platelet activating factor antagonist, and a neuroprotective agent (Maclennan et al., 2002). In contrast, PTX induces convulsions (Hasan et al., 2014). Yet, both block GABA and glycine receptors. Thus the mechanism of block may contribute to the opposing outcomes of these GABA/Gly antagonists. Alternatively, the selectivity of each antagonist for different receptor subtypes may explain its systemic physiology.

## Materials and Methods

### Cell Culture

Human embryonic kidney cells (HEK293 cells, American Type culture collection, MD) used in the experiments were cultured in 35 mm plates with Dulbecco’s Modified Eagle’s Medium (DMEM) complete with 10 % (v/v) Fetal Bovine Serum and 1% (v/v) penicillin-streptomycin. After 24 hours they were transfected using XtremeGENE 9 DNA Transfection reagent (Roche Inc, Basel Switzerland) with either α3 or α3+β cDNAs. GFP was co-transfected. A total of 2.5µg DNA were used per 35 mm dish with a ratio of 20:1 GlyR:GFP. When the cells were transfected with both α and β subunits, they were mixed in a 10β:1α ratio. Cells were cultured for 18-24 hours before recording.

### Electrophysiology

Whole cell patch clamp recordings were conducted on the transfected HEK 293 cells (visually identified by green fluorescence) using Axopatch 200B amplifier and pCLAMP 9.0 software (Axon Instruments, Foster City, CA). The drugs were applied through the Octaflow II (ALA Scientific Instruments, Inc. NY) localized superfusion system. Kreb’s external solution was used in recordings, containing (in mM): 140 NaCl, 4.7 KCl, 1.2 MgCl_2_, 2.5 CaCl_2_, 10 HEPES, 10 glucose (pH=7.4). Experiments were conducted using glass pipettes pulled by a multi-step horizontal puller (Sutter Instruments, CA) and fire polished. They had a tip resistance of ~5 MΩ and were filled with internal pipette solution containing (in mM): 140 CsMeSO_4_, 10 EGTA, 10 HEPES, 4 MgCl, 2 ATP-Mg (pH=7.2).

Glycine and picrotoxin was purchased from Sigma-Aldrich Corporation (St.Louis, MO) and Ginkgolide B from Tocris Biosciences (Bristol, UK). Glycine was dissolved in Kreb’s solution to make 5 mM solutions then further diluted to 500 µM. GB and PTX were dissolved in DMSO (Fisher Scientific, Waltham, MA) to make 10 mM stock solution and stored at 4 degree Celsius. The stock solutions were diluted with Krebs solution to the required concentrations before experiments.

## Results

### Glycine potency at the α3 GlyR

Initial experiments indicated that the α3 GlyRs, both homomeric and particularly heteromeric, are much less sensitive to glycine than GlyRs containing other alpha subunits. The glycine EC_50_ for α3 homomers is 166 μM, compared to 99 μM for α2 GlyRs (figure 1A, D). Furthermore, addition of the β subunit raised EC_50_ in α3β to 395 μM (figure 1B), while it had little effect in the α2 GlyR (figure 1C & D). In native tissue the glycine receptor is generally expressed as a heteromer and these results indicate that α3β GlyRs might be distinguished by their low glycine sensitivity (figure 1E).

**Figure 1:**
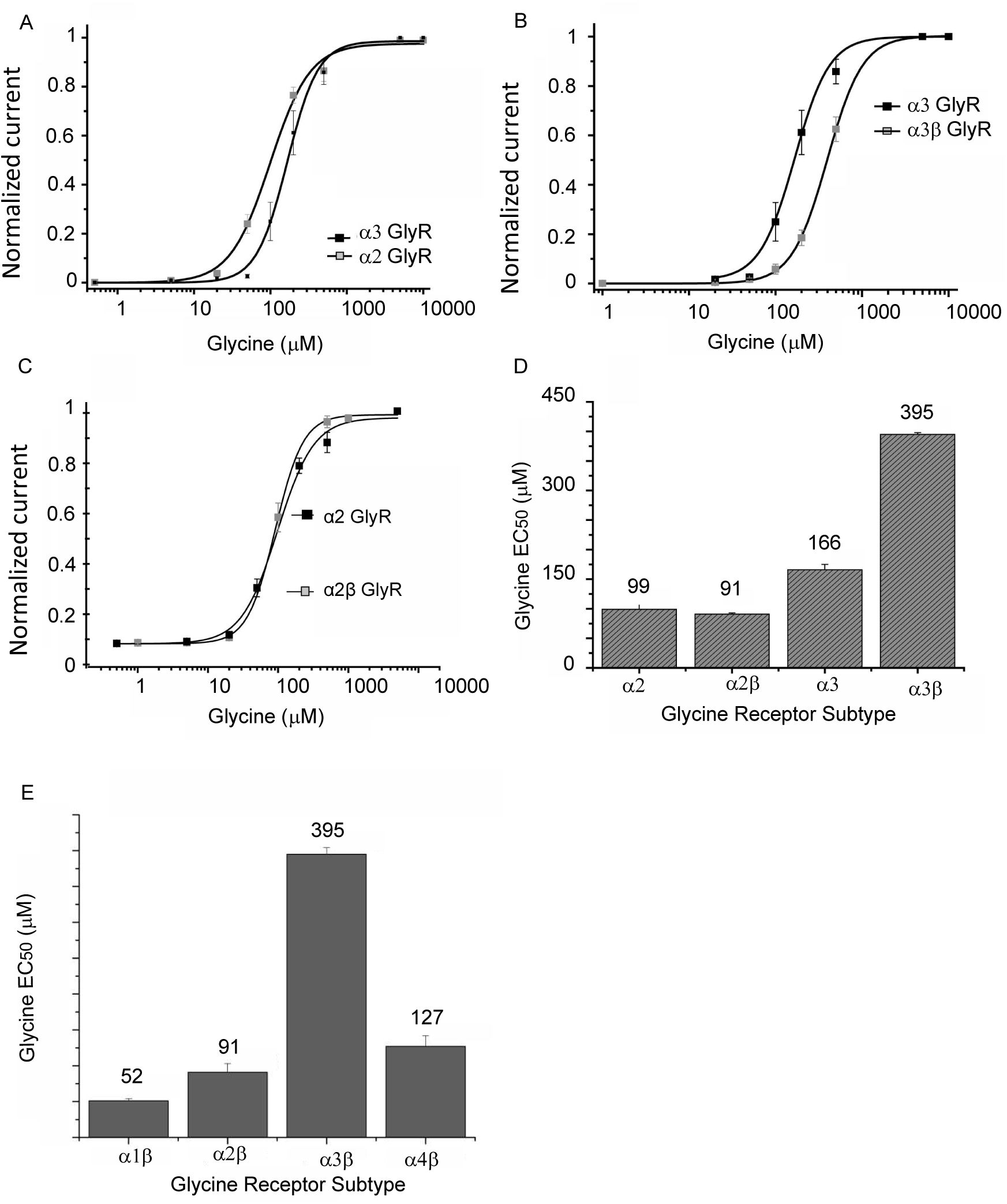
Comparative agonist sensitivity of the alpha subunits in glycine receptors. A) Relative response of homomeric alpha2 and alpha3 GlyRs. B) Glycine-elicited response in homomeric and heteromeric alpha3 GlyRs. C) Response of alpha2 homomeric and heteromeric GlyRs. D) EC50 values of homomeric and heteromeric alpha2 and alpha3 GlyRs. E) Comparison of mean EC50 values of various heteromeric GlyRs.

### Antagonist potency at the α3 GlyR

Two antagonists of ligand-gated chloride channels, picrotoxin and ginkgolide, have been used to compare GlyR subtypes. While α3 GlyRs are less sensitive to glycine than other subtypes, they have comparatively high sensitivity to picrotoxin. When tested against 200 μM glycine (approximate EC_60_), the picrotoxin IC_50_ was 0.9 μM. This compares with picrotoxin IC_50_ at α1 and α2 homomeric GlyRs of 37.3 and 12.6 μM, respectively (figure 2A). Like other GlyR subtypes, addition of the β subunit reduced the potency of picrotoxin (IC_50_ = 11.8 μM tested against the EC_60_ of glycine which is 468 μM, figure 2B).

**Figure 2.**
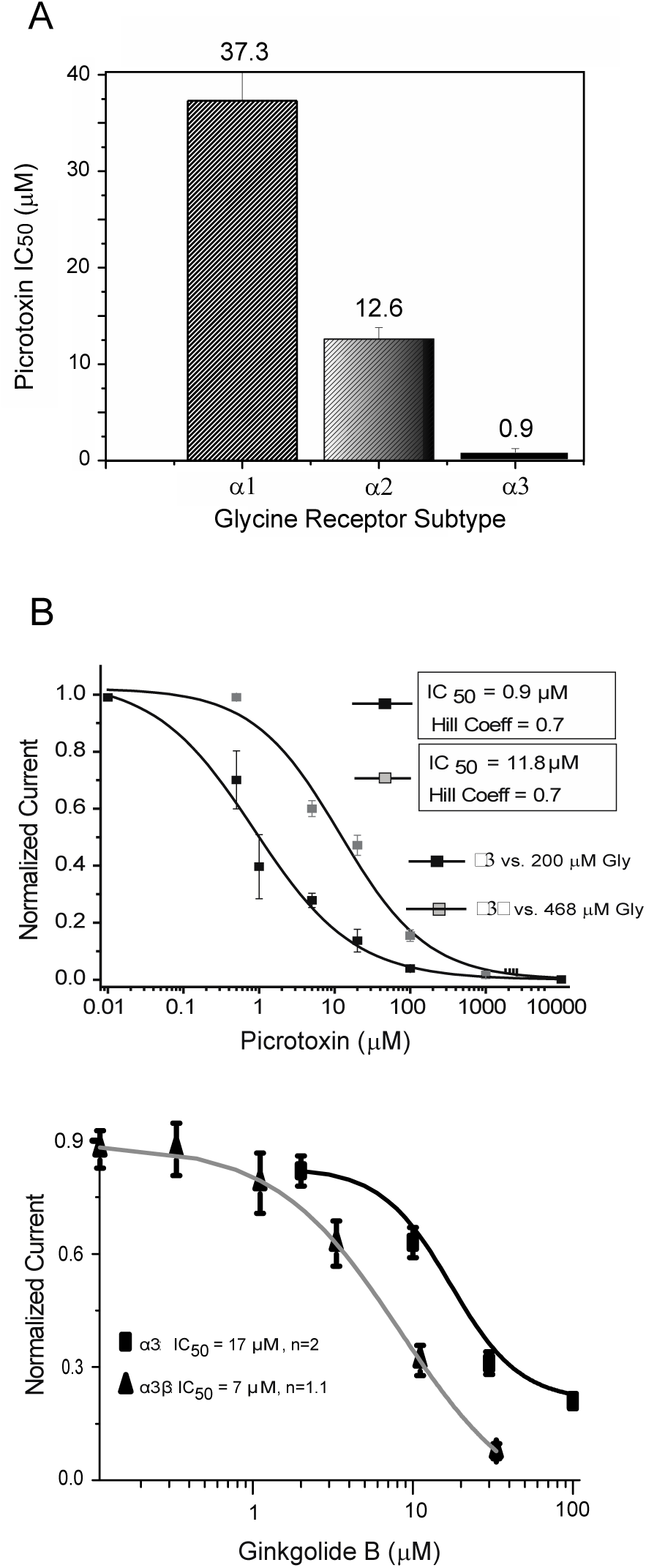
Antagonist effectiveness at GlyRs. A) Comparison of the IC50 values of picrotoxin block of glycine for homomeric GlyRs (see text for detailed description). B) Comparison of picrotoxin block of glycine activity in homomeric and heteromeric alpha3 GlyRs. C) Block of 500 uM glycine by ginkgolide in homomeric and heteromeric alpha3 GlyRs.

Ginkgolide B potency at the α3 GlyR was less than reported at α1 or α2 GlyRs, but like these other receptors the block was better in the heteromeric receptor (figure 2C). The GB IC_50_ is 17 μM in the α3 homomeric and 7 μM in the α3β heteromeric receptor. There is also a change in the Hill coefficient, from approximately 2 to 1. In comparison with α1 and α2 GlyRs, the relatively high potency of picrotoxin and low potency of ginkgolide B might be used to distinguish the α3β GlyR.

### GB is a non-competitive blocker and PTX is a competitive blocker of alpha 3 GlyRs

Studies have shown that GB is a non-competitive blocker of native GlyRs in hippocampal pyramidal rat neurons as well as in α1 and α2 GlyRs (Kondratskaya et al., 2004; Kondratskaya et al., 2005; Hawthorne et al., 2006a). In contrast, PTX is a competitive blocker in α1 and α2 GlyRs (Lynch et al., 1995; Wang et al., 2006). We tested GB and PTX block in α3 and α3β GlyRs.

The peak current response (I_0_) to application of 500 µM or 5 mM glycine was recorded and compared to the peak response in the presence of the antagonist (I), with either 2 µM PTX or 1 µM GB in cells held at −50 mV (figure 3). In 8 cells expressing homomeric α3 GlyR subunits, 1 µM GB on average blocked 12±9% of 500 µM glycine-induced current and 9±5% of 5 mM glycine-induced current (Figure 3A). There was no statistically significant difference between the two blocks, indicative of non-competitive inhibition. In contrast, 2 µM PTX blocked 43±9% of the 500 µM, but only 16±10% of 5 mM glycine-induced current. This difference in fractional block was statistically significant (p<0.01), suggesting that PTX is a competitive blocker.

**Figure 3:**
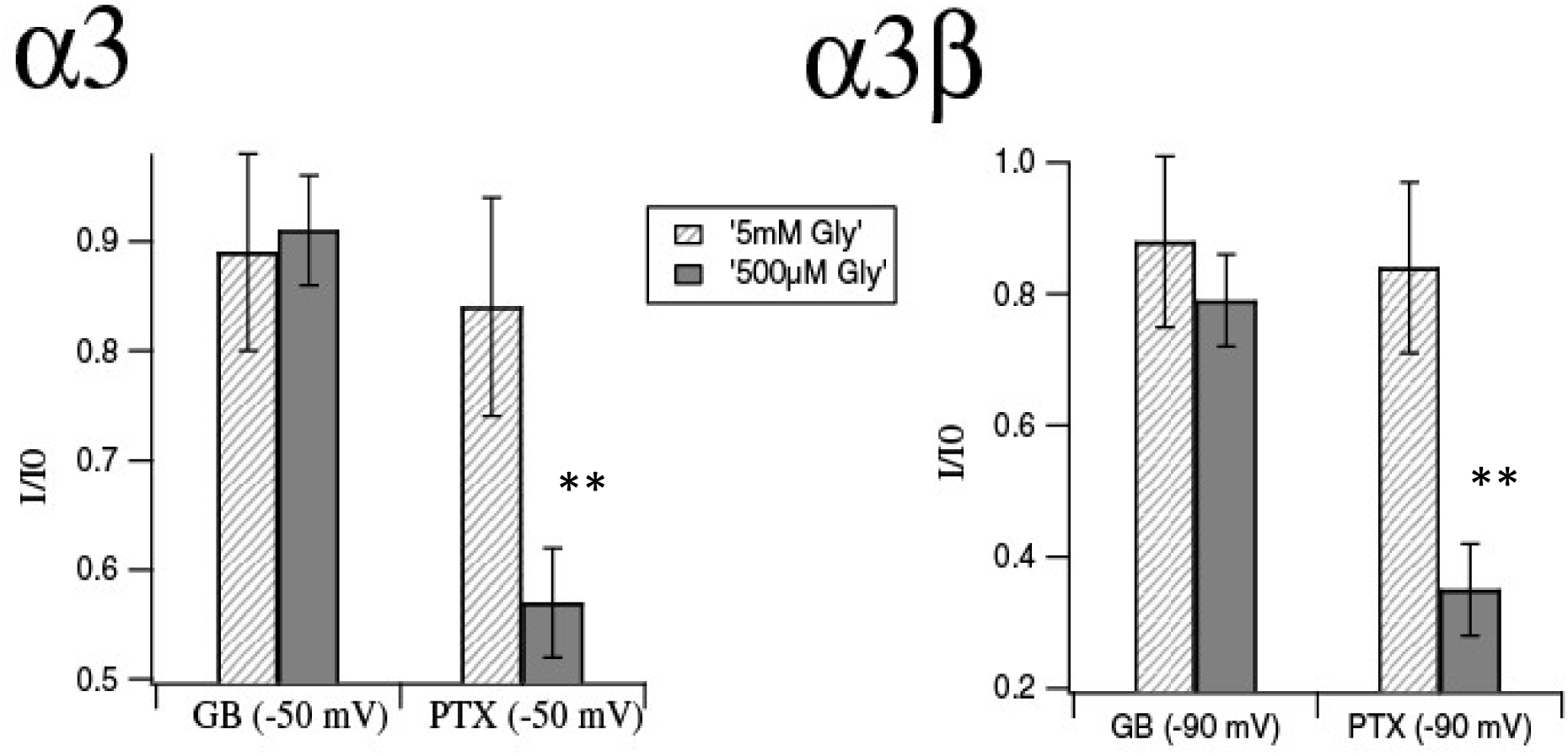
Picrotoxin is a competitive antagonist, GB is non-competitive. In the bar graphs, dashed bars represent I/I_0_ (where I is the peak glycine current in the presence of the antagonist and I_0_ is the peak current in glycine alone) for 5mM gly and solid bars represent I/I_0_ for 500 µM Gly. In α3 GlyR (left graph), blocking effects of 1 µM GB and 2 µM PTX were recorded at −50 mV. With GB, I/I_0_ (5 mM Gly) = 0.91±0.05, n=8 and I/I_0_ (500 µM Gly) = 0.88 ±0.09, n=8. With PTX, I/I_0_ (5 mM Gly) = 0.84±0.10, n=8 and I/I_0_ (500 µM Gly) = 0.57 ±0.09, n=8. The second graph shows the effect of 1 µM GB and10 µM PTX on α3β GlyR at −90 mV. With GB, I/I_0_ (5 mM Gly) = 0.88±0.07, n=8 and I/I_0_ (500 µM Gly) = 0.79 ±0.13, n=10. With PTX, I/I_0_ (5 mM Gly) = 0.84±0.10, n=5 and I/I_0_ (500 µM Gly) = 0.35 ±0.06, n=5.

In heteromeric α3β GlyR, there was no significant difference in the block of 500 µM or 5 mM glycine produced by 1 µM GB, but 10 µM PTX produced a greater block of 500 µM than 5 mM glycine (Figure 3B). 1 µM GB blocked 21±13% of 500 µM and 12±7% of 5 mM glycine-induced current. But 10 µM PTX blocked 65±6% of 500 µM and only 16±10% of 5 mM glycine-induced current. Higher PTX concentrations were used in the experiments on heteromeric receptors because they are less sensitive than the homomeric glycine receptor (see figure 2). Overall, block of α3-containing GlyRs by picrotoxin and ginkgolide B show the same competitive/non-competitive differences seen in the α1 and α2 containing GlyRs.

### GB block is voltage sensitive in alpha 3 GlyR

We tested the voltage sensitivity of GB and PTX block in homomeric and heteromeric α3 GlyR. Two measurements were taken during a 10 s application of 500 µM Glycine + antagonist: first the block of peak current and then after 5.5 sec. These responses were compared to the peak of the initial glycine current without antagonist (only the currents in the presence of antagonists are shown in figure 4). For homomeric α3 GlyRs, the measurements were taken at −10 mV and −50 mV, which were on either side of the reversal potential of −30 mV. A high concentration of GB (100 µM) was used in α3 GlyRs because of its comparatively low potency (Figure 4)

**Figure 4:**
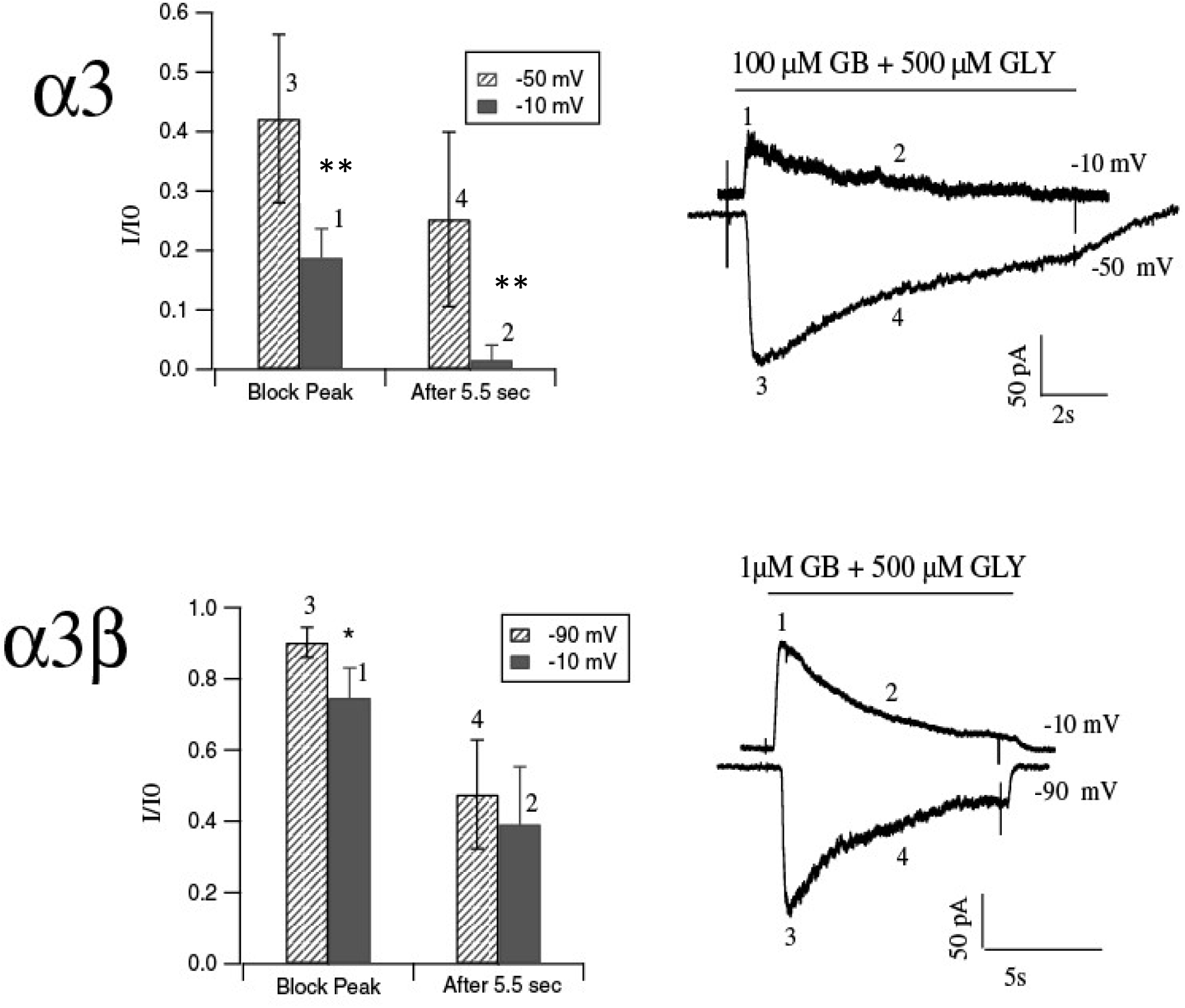
GB block is voltage-dependent in α3 GlyRs. Points marked 1,2,3 and 4 in the bar graph (left) correspond to the time points marked in the raw traces (right). The bar graph depicts I/I_0_ where I_0_ is the control 500 µM Glycine trace (not shown in raw trace). Top panel – Voltage sensitive GB block in α3 GlyRs – Points 1 and 3 are block of peak currents at −10 mV and −50 mV respectively. I/I_0_ values for the points marked 1 and 3 in the bar graph are 0.42±0.14, n=13 and 0.19±0.05, n=17. Points 2 and 4 are GB block after 5.5 sec at −10 mV and −50 mV respectively. I/I_0_ value for point 2 is 0.02±0.02, n=17 and for point 4 is 0.25±0.15, n=13. The peak of the control glycine current was same at −10 mV and −50 mV. Bottom Panel - Voltage sensitive GB block in α3β GlyRs - Points 1 and 3 are block of peak currents at −10 mV and −90 mV respectively. I/I_0_ value at point 1 is 0.75±0.08, n=9 and at point 3 is 0.90±0.04, n=7. Points 2 and 4 are GB block after 5.5 sec at −10 mV and −90 mV respectively. I/I_0_ value for point 2 is 0.39±0.16, n=9 and for point 4 is 0.47±0.15, n=7.

In α3 GlyRs the action of GB was voltage sensitive. GB blocked 58±14% of the glycine current at −50 mV in contrast to 81±5% at −10 mV. After 5.5 sec, GB blocked 75±15% at −50 mV and 98±2% at −10 mV (figure 4, top). Thus, at both time points α3 GlyRs were blocked significantly better at −10 mV compared to −50 mV (p<0.001). This finding is consistent with studies in rat hippocampal pyramidal neurons (Kondratskaya et al., 2002). Top right of Figure 4 shows sample traces of 10 s application of 500 µM Gly + 100 µM GB at the two voltages. The control glycine trace, without antagonist, is not shown because the control peak was of equal magnitude at both voltages. The histogram at top left summarizes the voltage sensitive block by GB of the peak current and the current after 5.5 seconds.

In α2, PTX is not a voltage sensitive blocker (Wang et al., 2006). In order to test how PTX behaves in α3, we used the same protocol employed for GB. At block peak, 2 µM PTX blocked 55±11% of the current evoked by 500 µM Gly at −10 mV and 52±13% at −50 mV. Since the differences in blocks are not significant, we conclude that PTX is not a voltage sensitive blocker in α3 GlyRs (data not shown).

Similar experiments were repeated on α3β GlyRs, but using a lower GB (1 µM) concentration because of its greater potency in the heteromers. Here we made the measurements at −10 mV, −50mV and −90 mV. At −50mV and −10 mV this batch of cells showed outward current, so we also tested at −90 mV, where an inward current was evoked.

In α3β GlyRs, GB blocked 25±8% of 500 µM Gly induced peak current at −10 mV, compared to 10±4% at −90 mV and 11±4% at −50 mV. Thus, like homomeric α3 GlyRs, there was a significantly better GB block at −10 mV in α3β GlyRs. Since there was almost no difference between −90 mV and −50 mV block, we conclude that direction of ion flow is not an important factor in GB block. There was not a statistically significant difference in block at the two voltages after 5.5 s. GB block of glycine in α3β GlyRs was 61±16% at −10 mV, 53±15% at −90 mV, and 50±16% at −50 mV. The right panel of Figure 4B shows sample traces and the left panel shows a summary of the data at −10 mV and −90 mV. Difference in the rate of desensitization in α3β and α3 GlyRs is a potential reason for the lack of voltage sensitivity after 5.5 s in α3β.

The α3β GlyRs showed no statistically significant difference in PTX block at −10 mV, −50 mV and −90 mV. At the peak of the glycine current, PTX blocked 30±13% at −10 mV, 30±13% at −50mV and 38±15% at −90 mV.

### Differences from other GlyRs

a. Pre-treatment with GB and PTX blocks α3 GlyR currents

Both GB and PTX are thought to block the channel pore. Many pore blockers are use-dependent because they require the agonist to open the pore to allow access to the blocker. Hence, two blocking mechanisms were compared: 1) Liganded or use-dependent block that occurs when glycine and antagonist are applied together and 2) Unliganded or non-use-dependent block in which the action of the antagonist does not depend on the presence of the agonist. PTX block has been linked to amino acids in the glycine channel pore but has been shown to be non-use dependent in α1 (Lynch et al., 1995), but use-dependent in α2 (Wang et al., 2006). GB has been proposed to block in a use-dependent manner in native hippocampal neuron (Kondratskaya et al., 2002) and in α2 GlyRs (Hawthorne et al., 2006a). In our experiments, the protocol used to test for unliganded block was to: 1) apply 500 µM glycine for 2 s (first glycine response), wait for the current to return to baseline, and then 2) apply antagonist GB or PTX (without glycine) for 10 s and then after a wash period of 3 s, 3) apply only 500 µM glycine for 2 s (second glycine response). If the antagonist blocked the unliganded receptor, then the second glycine peak would be reduced. To test for liganded block: 1) 500 µM glycine was first applied for 2 s (first glycine response) then after recovery to baseline 2) GB or PTX was co-applied with 500 µM glycine for 10 s, followed by a 3 s wash and then 3) 500 µM glycine (without antagonist) was reapplied for 2 s (second glycine peak). The results are expressed as the percentage of control peak glycine current that was blocked 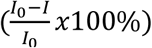.

An example of this protocol is shown in Figure 5. The experiment was performed on α3 homomers and PTX fully blocked the effect of concomitantly applied glycine (top trace). After washout of both drugs, brief reapplication of glycine caused progressively larger currents, indicating a slow removal of the bound PTX block. When this protocol was repeated with the application of PTX alone (bottom trace) there was a similar slow unblocking. Thus, regardless of whether PTX was applied in the presence (liganded) or absence (unliganded) of glycine, PTX produced a similar level of block. Each of the secondary glycine applications represents a new sequence of the three steps and all sequences shown were performed in the same cell. The results indicate that, like the α1 and α2 GlyRs, PTX block of the α3 homomer was not use-dependent.

**Figure 5:**
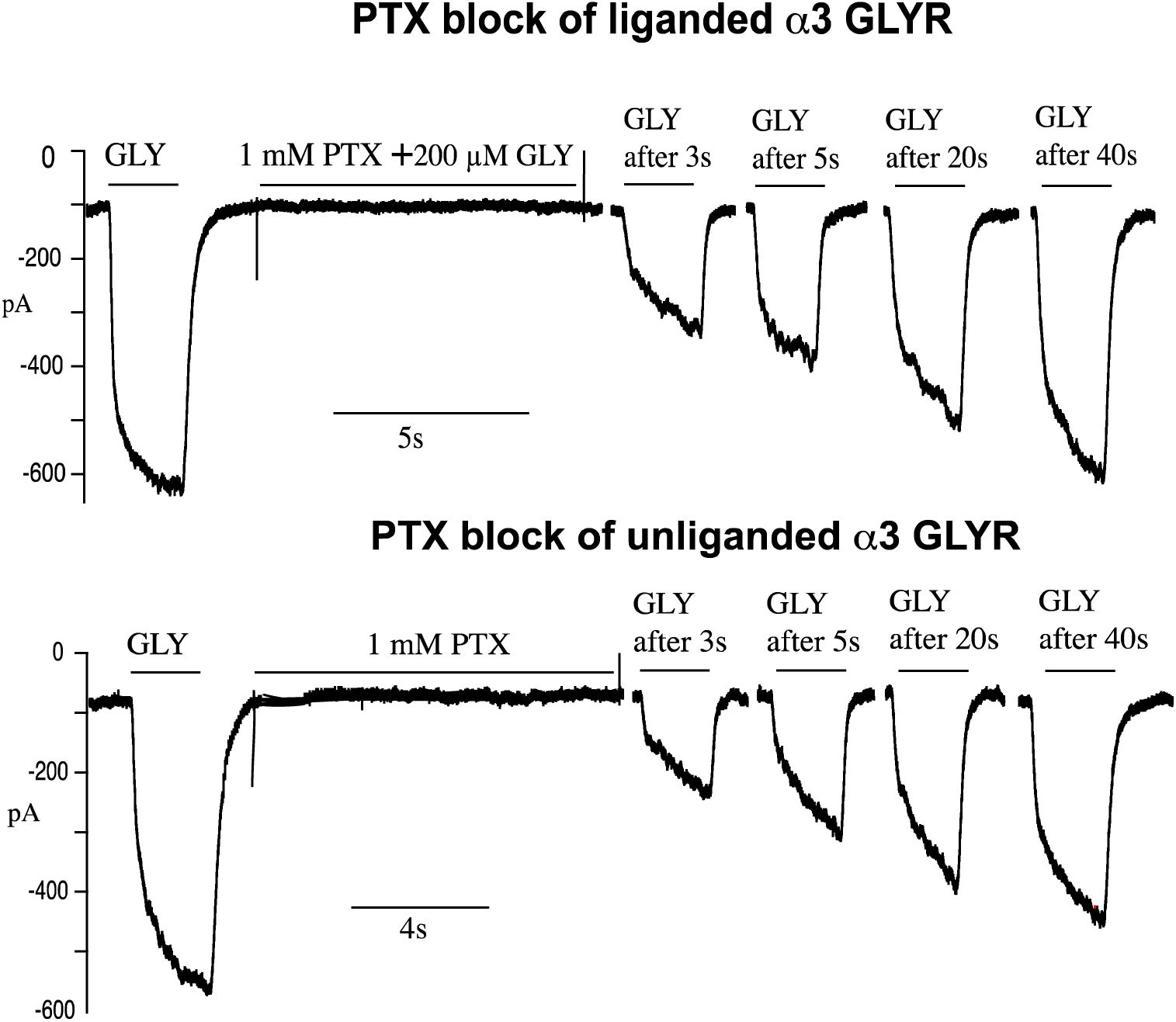
Picrotoxin block of liganded and unliganded homomeric alpha3 glycine receptors. Glycine was applied before and then again after treatment with picrotoxin plus glycine (top) or picrotoxin alone (bottom). Each of the four post-picrotoxin traces represents a separate experiment in which glycine was applied sometime after picrotoxin removal, as indicated.

To test the validity of our experimental protocol, we compared the GB liganded and unliganded block of α2 GlyRs, using the protocol shown in figure 5 (Figure 6, top left). The first pair of superimposed traces shows that glycine alone (black) and glycine after pretreatment with GB (gray) produced similar currents. There was no evidence of α2 GlyR inhibition in the unliganded-block protocol. In contrast, ~27% of α2 GlyRs were blocked in liganded receptors. This experiment agrees with studies indicating that α2 GlyRs show only liganded block and also supports our experimental design.

**Figure 6:**
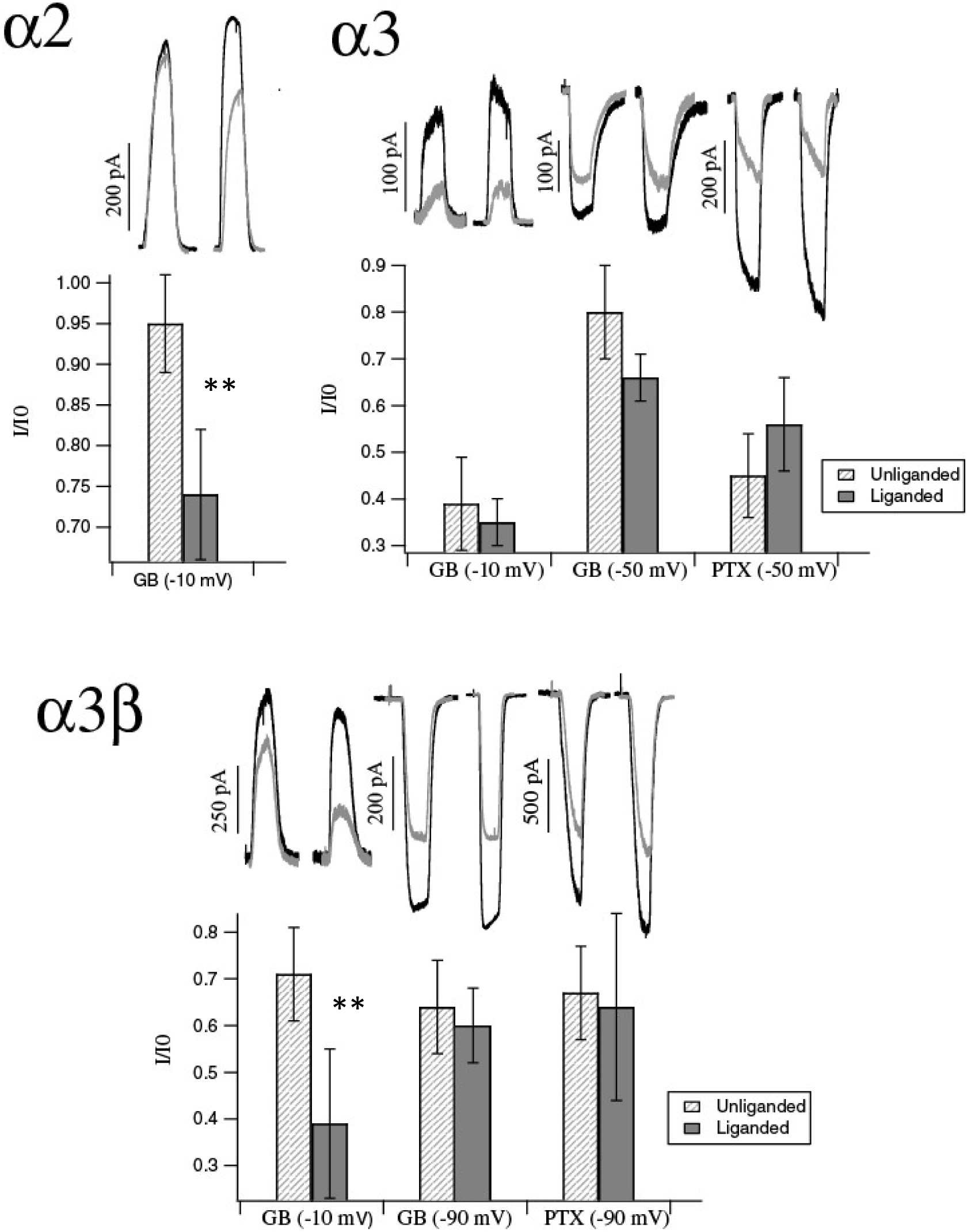
GB can block the unliganded glycine receptor. The figure shows blocking effects of antagonists GB and PTX in liganded and unliganded state on α2 α3 and α3β GlyRs. For all parts dashed bars represent unliganded block conditions while solid bar represents liganded block conditions. The raw traces above dashed bars show relative peak of the first glycine peak (black) to second glycine peak after 3s wash (grey). The bars represent I/I_0_ values where I_0_ is the first glycine peak and I is the second glycine peak. Error bars represent SD. In α2 part of the figure, I/I_0_ value for liganded state at −10 mV is 0.74 ± 0.08, n=12. The unliganded block was fully recovered by 3s (I/I_0_=0.95± 0.06, n=15). In α3 GlyRs, I/I_0_ values at −10 mV GB unliganded was 0.39±0.1, n=5 and at −10 mV GB liganded was 0.35±0.05, n=5. The plotted value at −50 mV GB unliganded was 0.80± 0.1, n=3 and liganded was 0.66 ± 0.05, n=4. For PTX unliganded block, I/I_0_ value was 0.45±0.09, n=8 and for PTX liganded block it was 0.56±0.1, n=7. In α3β GlyRs, I/I_0_ values at −10 mV GB unliganded was 0.71±0.1, n=8 and for liganded GB at the same voltage it was 0.39 ± 0.16, n=11. At −90 mV, plotted I/I_0_ values for unliganded GB was 0.64±0.1, n=8 and liganded GB was 0.60±0.08, n=7. For PTX block, I/I_0_ values at unliganded was 0.67±0.1, n=6 and at liganded it was 0.64±0.20, n=6.

Surprisingly, GB blocked unliganded α3 GlyRs and this block showed voltage dependence similar to the liganded receptor block (Figure 6, top right). Each superimposed pair of current traces in Figure 6 shows the response to glycine alone in black and the second glycine current, 3s after GB removal, in gray. The bar graph below each pair of superimposed traces shows the fractional remaining current in the step 3 protocol for all cells tested. The first two sets of superimposed traces in Figure 6 top right show that, for cells held at −10 mV, GB blocks on average 61±11% of the unliganded receptor after 3s and 65±5% of the liganded receptor. The middle two traces show that GB is less effective in cells held at −50 mV, where the unliganded block is 20±5% and the liganded block is 34±5%. Thus, GB could bind and block the closed channel. PTX could also bind to the closed, unliganded α3 GlyR as shown in the last two pair of superimposed traces. In the unliganded protocol, 55±9% of the current remained blocked by PTX after 3s, (measurements only conducted at −50 mV since PTX block is not voltage sensitive). PTX blocked 44±11% of the GlyR current in the liganded receptor. In all three sets of responses there was no statistically significant difference in the fraction of block produced in the liganded vs the unliganded state.

These experiments were repeated in the heteromeric GlyR, but drug concentrations were 500 µM glycine, 1µM GB and 10 µM PTX, because the heteromeric receptor was more sensitive to GB and less to PTX and glycine (Figure 6, lower panel). These receptors were also found to be blocked in unliganded and liganded state by PTX and GB. Again, we tested α3β GlyRs at −10 mV, −50 mV (not shown) and −90 mV to examine whether direction of ion flow has any effect on recovery. At −10 mV and −50 mV the currents were outward and at −90 mV the current was inward. For the unliganded receptor, the remaining GB block after 3s was 29±13% at −10 mV, 40±15% at −50 mV and 36±14% at −90 mV. When these receptors were tested in the liganded state, the GB block was 61±16% at −10mV and 40±9% at both −90 mV and at −50 mV. The liganded PTX block after 3s was 36±11%, measured at −90 mV holding potential.

The α3 homomeric and heteromeric glycine receptors are blocked in the unliganded and liganded state by GB and PTX. Only −10 mV and −90 mV GB blocks are shown in figure 7 because there were no differences between GB blocks at −50 mV vs −90 mV. Thus, it follows that direction of ion flow does not affect recovery. Based on these results, the next set of experiments were conducted at −10 mV and at −90 mV.

**Figure 7:**
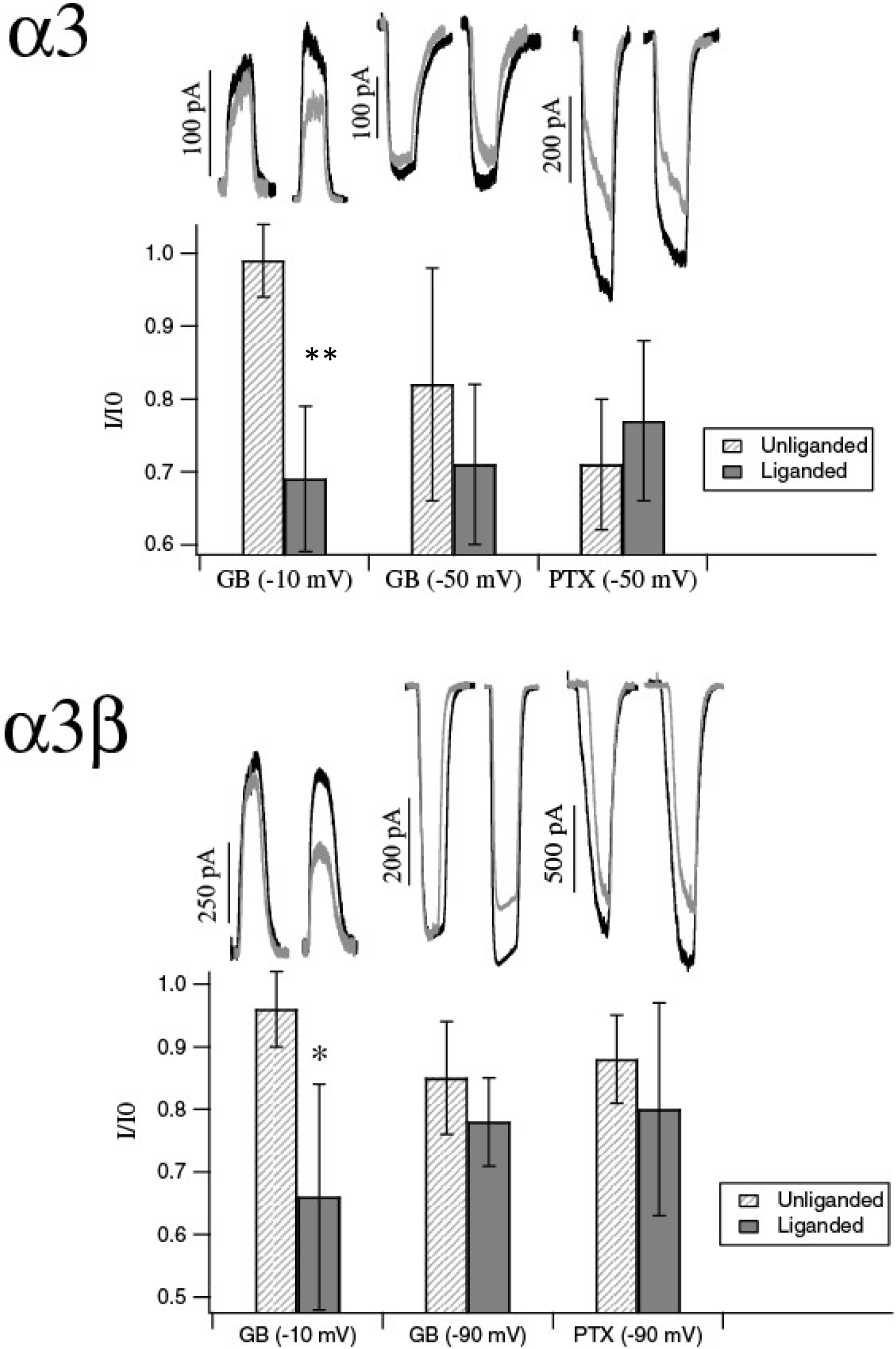
Recovery from GB block of glycine receptors is voltage dependent. Shows recovery from unliganded and liganded blocks of GB and PTX after 10s wash. For both α3 and α3β GlyRs dashed bars and the corresponding traces above that represent unliganded block while solid bars along with the raw traces above represent liganded block. In the raw traces, black shows the first glycine response and grey shows the glycine response after 10s wash. The plotted values in the bar graph is I/I_0_ values where I_0_ is the first glycine peak and I is the second glycine peak. Error bars represent SD. α3 GlyRs – Plotted I/I_0_ values for unliganded GB block at −10 mV = 0.99±0.05, n=5 and liganded GB block = 0.70±0.10, n=5 (p<0.05). At −50 mV, unliganded GB block was 0.83±0.16, n=3 and liganded GB block was 0.71±0.10, n=4. With unliganded PTX block I/I_0_ value was 0.72±0.09, n=7 and with liganded PTX I/I_0_ was 0.77±0.11, n=8. α3β GlyRs - At −10 mV, I/I_0_ values for unliganded GB block was 0.96±0.08, n=6 and liganded GB block was 0.66±0.07, n=6 p<0.001. At −90 mV, I/I_0_ values for unliganded GB block was 0.85±0.09, n=7 and liganded GB block was 0.78±0.07, n=6. I/I_0_ values for unliganded PTX block was 0.88±0.07, n=6 and liganded PTX block was 0.80±0.16, n=6.

4. Recovery Rate from Liganded and Unliganded block

Next, we compared the rate of recovery of the GlyRs from the liganded and unliganded block described above. The same drug concentrations were used (i.e, 500 µM Glycine, 100 µM GB, 1 mM PTX for α3 and 500 µM Glycine, 1 µM GB and 10 µM PTX for α3β) but the wash time was prolonged to 10 s. The results express GB or PTX block remaining after 10s wash. Note that the peak of the second glycine current after 10 sec includes recovery from both block and desensitization. Preliminary experiments, without antagonists, indicated that recovery from desensitization was complete at 3s. However, it is unclear if antagonists will affect this rate of recovery from desensitization, so the recovered current of the second glycine peak may not only measure of antagonist unbinding.

At −10 mV, homomeric α3 GlyRs recovered faster from GB unliganded block compared to GB liganded block (Figure 8A). There was a 30±12% block remaining in the liganded condition after 10s and the receptors fully recovered in ~60 s. Unliganded GB block fully recovered in 10 sec. There were no significant difference in recovery after 10s of liganded and unliganded blocks with GB at −50 mV or for PTX at −50 mV. After 10 s wash, at −50 mV, about 17±17% block remained in unliganded GB condition and about 28±11% block remained in liganded GB condition. Similarly, block remaining 10 s after unliganded PTX treatment was 28±9% and after liganded PTX was 23±11%.

A similar trend was found in α3β heteromers. As shown in Figure 8B, liganded GB block at −10 mV produced a 34±17% block after 10s wash while only 5±5% block remained 10s after unliganded GB block at −10mV. No significant difference between recovery from liganded and unliganded block was found with GB at −90 mV or with PTX. At −90 mV, block remaining 10s after liganded GB block was 15±7% and after unliganded GB block was 22±9%. And 20±17% block remained 10s after liganded PTX block and 12±7% block remained after unliganded PTX block.

These results suggest mechanistic differences between GB block of the glycine bound and glycine free receptor. It is possible that there is more than one voltage sensitive binding site for GB. For simplicity if we assume there are two sites, one site is accessible in the liganded or unliganded state and the other site is only accessible when glycine is present. At holding potential of −10 mV, GB is bound efficiently to the second site and dissociates more slowly compared to the same liganded receptors at more negative voltages or to unliganded receptors at −10 mV.

## Discussion

### Distinct Pharmacology of α3 GlyR

This study reveals a number of unique features conferred by the α3 subunit of the glycine receptor. It is relatively insensitive to glycine, the heteromeric α3β receptor EC_50_ is close to 400 μM. This is almost an order of magnitude less sensitive than heteromeric α1 GlyRs. The β subunit contributes to this low glycine sensitivity, which is unusual in that the β subunit of α1 or α2 GlyRs has little effect on sensitivity. The low glycine sensitivity might expand the dynamic range of the glycine response, since the α3-containing receptor can respond to glycine levels that have saturated the other GlyR types.

Another interesting property of the α3 GlyR is the high sensitivity to picrotoxin. With an IC_50_ of ~1 μM, the homomeric α3 GlyR is as sensitive, or more sensitive, than GABA receptors. The α3β GlyR has an PTX IC_50_ of ~12 μM, which is still close to that of GABA receptors. This indicates that PTX does not uniquely identify GABA receptors in pharmacological experiments and adds to the questionable utility of PTX based on studies of α2 GlyRs.

### PTX Block of α3 GlyR

Mechanism of PTX inhibition in ligand-gated ion channels has not been fully elucidated. A single mutation in M2 reduces PTX sensitivity in GABA and glycine receptors (Pribilla et al., 1992; Zhang et al., 1995). The ring of threonines in the 6’ position is associated with PTX sensitivity in GABA, 5-HT3 (Gurley et al., 1995; Das and Dillon, 2005) and glycine receptors (Shan et al., 2001a; Hawthorne and Lynch, 2005), which suggests a pore binding model for PTX. However, PTX inhibition is competitive and voltage insensitive (Lynch et al., 1995; Wang et al., 2006) which are inconsistent with a pore blocker. This has led to an allosteric model of PTX inhibition (Lynch et al., 1995; Shan et al., 2001a). PTX is competitive, not use-dependent, and voltage insensitive in α3 GlyRs. Furthermore, recovery rate of α3 GlyR from liganded and unliganded block by PTX is indistinguishable. Thus, PTX binding site in α3 GlyRs is in a region that is freely accessible in both open and closed state of the receptor. Glycine is not necessary for the α3 GlyR to recover from liganded block, which argues against the trap mechanism that is proposed in α2 GlyRs (Wang et al., 2006) and α1 GlyRs (Hawthorne and Lynch, 2005).

### Comparison of PTX and GB at α3 GlyR

PTX and GB have similarities in molecular structure (Ivic et al., 2003a) and are both thought to bind near the pore region of the channel (Hawthorne et al., 2006b; Heads et al., 2008b). Support for this comes from mutations of T6, an amino acid in the middle of the M2 transmembrane segment, that reduce the effectiveness of PTX and GB (Shan et al., 2001b; Hawthorne et al., 2006b). In our experiments the potency of GB was similar to PTX, although repeated exposure can increase the apparent potency of a use-dependent ligand. The heteromer is more sensitive to GB and less sensitive to PTX, which is also true in α2β GlyRs. The most distinctive differences between the two antagonists is that GB is voltage-dependent and non-competitive. These are properties that the α3 GlyR shares with the α2 receptor. In heterologously expressed α2 GlyRs, and in native rat hippocampal neurons, GB potency does not depend on the glycine concentration but is less effective when the cell is hyperpolarized. These two properties, along with use-dependence, are hallmarks of pore blockers.

But GB block at α3 GlyRs is not exclusively use-dependent; GB can bind and block the unliganded, closed state of the receptor. This is unlike GB’s effect in hippocampal pyramidal cells or in heterologously expressed α2 GlyRs and does not fit with the classic pore blocker model which acts only on open channels.

Although α1 GlyRs has been reported to be blocked in unliganded state by PTX, none of the other receptors are blocked by either antagonist in the unliganded state. The unique antagonism profile of α3 GlyRs may help in pharmacologically identifying the subunit composition in native neurons.

Most studies agree that homomeric α3 GlyRs are not expressed in somatic regions of adult neurons (Lynch, 2009), but may be present in presynaptic nerve terminals of adult auditory brainstem neurons (Turecek and Trussell, 2002) and neurons of rat supraoptic nucleus (Deleuze et al., 2005).The α3β GlyRs were seen in the inner plexiform layer of the retina and dorsal horn of the spinal cord neurons using immunohistochemistry (Haverkamp et al., 2003; Harvey et al., 2004), but these findings need validation (Elliott et al., 2015). Use of GB and PTX as pharmacological tools can help provide further support for this identification. α3 GlyRs have been implicated in tactile allodynia (Huang et al., 2017) and Glra3^-/-^ knockout mice have reduced prostaglandin E2 induced pain sensitization. (Harvey et al., 2004). These findings identify α3 GlyRs as molecular pain target and since GB and PTX block α3β GlyRs, they might be useful in pain therapy if their action can be localized.

We sought to compare PTX antagonism with GB antagonism in α3 GlyRs due to the postulated structural similarity between the two compounds (Ivic et al., 2003b). Although the mechanism of GB inhibition is unresolved, there is evidence that GB binds to the 2’-6’ pore lining region of the M2 transmembrane segment of GlyRs (Kondratskaya et al., 2005; Hawthorne et al., 2006a; Heads et al., 2008a). This is further supported by the non-competitive, use-dependent and voltage sensitive nature of the block (Kondratskaya et al., 2002; Kondratskaya et al., 2004; Kondratskaya et al., 2005; Hawthorne et al., 2006a). In addition to non-competitive, voltage sensitive and use-dependent properties, our data show an unliganded GB block in α3 GlyRs. We have also shown that recovery from the liganded block is slower than the unliganded block. These data might be explained by two GB binding sites in the α3 GlyR. Site 1 is outside the pore and responsible for unliganded block. It can even be in the extracellular domain since a recent study identifies a novel allosteric site above the glycine binding site (Huang et al., 2004). Site 2 is in the pore and accessible only when the channel is open. Recovery from liganded GB block requires GB to fall off from both Sites 1 and 2 and hence takes longer than unliganded recovery. Either or both of these sites can be voltage sensitive. A similar competitive, unliganded inhibition of ρ1 GABA_C_ receptors by GB has been reported (Huang et al., 2012) and they also suggested the possibility of two ginkgolide binding sites. The apparent competitive inhibition in GABA_C_ receptors may be due to an allosteric effect on agonist binding rather than a physical competition for the binding site.

## Acknowledgement

The work was funded by National Eye Institute grant to M.M.S. S.C. was supported by scholarships from University at Buffalo.

## References

Baer K, Waldvogel Hj Fau - Faull RLM, Faull RI Fau - Rees MI, Rees MI (2009) Localization of glycine receptors in the human forebrain, brainstem, and cervical spinal cord: an immunohistochemical review.

Crook J, Hendrickson A, Robinson FR (2006) Co-localization of glycine and gaba immunoreactivity in interneurons in Macaca monkey cerebellar cortex. Neuroscience 141:1951-1959.

Das P, Dillon GH (2005) Molecular determinants of picrotoxin inhibition of 5-hydroxytryptamine type 3 receptors.

Deleuze C, Runquist M Fau - Orcel H, Orcel H Fau - Rabie A, Rabie A Fau - Dayanithi G, Dayanithi G Fau - Alonso G, Alonso G Fau - Hussy N, Hussy N (2005) Structural difference between heteromeric somatic and homomeric axonal glycine receptors in the hypothalamo-neurohypophysial system.

Durisic N, Godin AG, Wever CM, Heyes CD, Lakadamyali M, Dent JA (2012) Stoichiometry of the Human Glycine Receptor Revealed by Direct Subunit Counting. The Journal of Neuroscience 32:12915-12920.

Elliott K, McQuaid S, Salto-Tellez M, Maxwell P (2015) Immunohistochemistry should undergo robust validation equivalent to that of molecular diagnostics.

Gurley D, Amin J Fau - Ross PC, Ross Pc Fau - Weiss DS, Weiss Ds Fau - White G, White G (1995) Point mutations in the M2 region of the alpha, beta, or gamma subunit of the GABAA channel that abolish block by picrotoxin.

Harvey RJ, Depner UB, Wässle H, Ahmadi S, Heindl C, Reinold H, Smart TG, Harvey K, Schütz B, Abo-Salem OM, Zimmer A, Poisbeau P, Welzl H, Wolfer DP, Betz H, Zeilhofer HU, Müller U (2004) GlyR α3: An Essential Target for Spinal PGE2-Mediated Inflammatory Pain Sensitization. Science 304:884-887.

Hasan ZA, Abdel Razzak RL, Alzoubi KH (2014) Comparison between the effect of propofol and midazolam on picrotoxin-induced convulsions in rat. Physiology & Behavior 128:114-118.

Haverkamp S, Muller U Fau - Harvey K, Harvey K Fau - Harvey RJ, Harvey Rj Fau - Betz H, Betz H Fau - Wassle H, Wassle H (2003) Diversity of glycine receptors in the mouse retina: localization of the alpha3 subunit.

Hawthorne R, Lynch JW (2005) A picrotoxin-specific conformational change in the glycine receptor M2-M3 loop.

Hawthorne R, Cromer BA, Ng H-L, Parker MW, Lynch JW (2006a) Molecular determinants of ginkgolide binding in the glycine receptor pore. Journal of Neurochemistry 98:395-407.

Hawthorne R, Cromer BA, Ng HL, Parker MW, Lynch JW (2006b) Molecular determinants of ginkgolide binding in the glycine receptor pore. J Neurochem 98:395-407.

Heads JA, Hawthorne RL, Lynagh T, Lynch JW (2008a) Structure-activity analysis of ginkgolide binding in the glycine receptor pore. Journal of Neurochemistry 105:1418-1427.

Heads JA, Hawthorne RL, Lynagh T, Lynch JW (2008b) Structure-activity analysis of ginkgolide binding in the glycine receptor pore. J Neurochem 105:1418-1427.

Heinze L, Harvey RJ, Haverkamp S, Wässle H (2007) Diversity of glycine receptors in the mouse retina: Localization of the α4 subunit. The Journal of Comparative Neurology 500:693-707.

Huang SH, Duke RK, Chebib M, Sasaki K, Wada K, Johnston GAR (2004) Ginkgolides, diterpene trilactones of Ginkgo biloba, as antagonists at recombinant α1β2γ2L GABAA receptors. European Journal of Pharmacology 494:131-138.

Huang SH, Lewis TM, Lummis SC, Thompson AJ, Chebib M, Johnston GA, Duke RK (2012) Mixed antagonistic effects of the ginkgolides at recombinant human rho1 GABAC receptors. Neuropharmacology 63:1127-1139.

Huang X, Shaffer PL, Ayube S, Bregman H, Chen H, Lehto SG, Luther JA, Matson DJ, McDonough SI, Michelsen K, Plant MH, Schneider S, Simard JR, Teffera Y, Yi S, Zhang M, DiMauro EF, Gingras J (2017) Crystal structures of human glycine receptor [alpha]3 bound to a novel class of analgesic potentiators. Nat Struct Mol Biol 24:108-113.

Ivic L, Sands TT, Fishkin N, Nakanishi K, Kriegstein AR, Stromgaard K (2003a) Terpene trilactones from Ginkgo biloba are antagonists of cortical glycine and GABA(A) receptors. J Biol Chem 278:49279-49285.

Ivic L, Sands Tt Fau - Fishkin N, Fishkin N Fau - Nakanishi K, Nakanishi K Fau - Kriegstein AR, Kriegstein Ar Fau - Stromgaard K, Stromgaard K (2003b) Terpene trilactones from Ginkgo biloba are antagonists of cortical glycine and GABA(A) receptors.

Kondratskaya EL, Lishko PV, Chatterjee SS, Krishtal OA (2002) BN52021, a platelet activating factor antagonist, is a selective blocker of glycine-gated chloride channel. Neurochemistry International 40:647-653.

Kondratskaya EL, Fisyunov AI, Chatterjee SS, Krishtal OA (2004) Ginkgolide B preferentially blocks chloride channels formed by heteromeric glycine receptors in hippocampal pyramidal neurons of rat. Brain Research Bulletin 63:309-314.

Kondratskaya EL, Betz H, Krishtal OA, Laube B (2005) The β subunit increases the ginkgolide B sensitivity of inhibitory glycine receptors. Neuropharmacology 49:945-951.

Lynch JW (2009) Native glycine receptor subtypes and their physiological roles. Neuropharmacology 56:303-309.

Lynch JW, Rajendra S Fau - Barry PH, Barry Ph Fau - Schofield PR, Schofield PR (1995) Mutations affecting the glycine receptor agonist transduction mechanism convert the competitive antagonist, picrotoxin, into an allosteric potentiator.

Maclennan KM, Darlington CL, Smith PF (2002) The CNS effects of Ginkgo biloba extracts and ginkgolide B. Progress in Neurobiology 67:235-257.

Pribilla I, Takagi T, Langosch D, Bormann J, Betz H (1992) The atypical M2 segment of the beta subunit confers picrotoxinin resistance to inhibitory glycine receptor channels. The EMBO Journal 11:4305-4311.

Shan Q, Haddrill Jl Fau - Lynch JW, Lynch JW (2001a) A single beta subunit M2 domain residue controls the picrotoxin sensitivity of alphabeta heteromeric glycine receptor chloride channels.

Shan Q, Haddrill JL, Lynch JW (2001b) A single beta subunit M2 domain residue controls the picrotoxin sensitivity of alphabeta heteromeric glycine receptor chloride channels. J Neurochem 76:1109-1120.

Turecek R, Trussell LO (2002) Reciprocal developmental regulation of presynaptic ionotropic receptors.

Wang D-S, Mangin J-M, Moonen G, Rigo J-M, Legendre P (2006) Mechanisms for Picrotoxin Block of α2 Homomeric Glycine Receptors. Journal of Biological Chemistry 281:3841-3855.

Xu T-L, Gong N (2010) Glycine and glycine receptor signaling in hippocampal neurons: Diversity, function and regulation. Progress in Neurobiology 91:349-361.

Zhang D, Pan Zh Fau - Zhang X, Zhang X Fau - Brideau AD, Brideau Ad Fau - Lipton SA, Lipton SA (1995) Cloning of a gamma-aminobutyric acid type C receptor subunit in rat retina with a methionine residue critical for picrotoxinin channel block.

